# Reconfiguration of tumor cells with LCOR transcription factor mRNA nanotherapy to enhance immunotherapy efficacy

**DOI:** 10.1101/2025.04.02.646434

**Authors:** Gabriel Serra-Mir, Elena Haro-Martínez, Chiara Cannatà, José Ángel Palomeque, Sandra Blasco-Benito, Anna López-Plana, María Sanz-Flores, Marc Sabartés, Cristina Fornaguera, Toni Celià-Terrassa

**Affiliations:** Cancer Research Program, Hospital del Mar Research Institute (HMRI), Barcelona, Spain; Grup d’Enginyeria de Materials (Gemat), Institut Químic de Sarrià (IQS), Universitat Ramon Llull (URL), Barcelona, Spain; Institució Catalana de Recerca i Estudis Avançats (ICREA), Barcelona, Spain; Centro de Investigación Biomédica en Red de Oncología (CIBERONC-ISCIII), Madrid, Spain

**Keywords:** Immunotherapy, mRNA therapy, transcription factor, tumor cell reprogramming, triple negative breast cancer

## Abstract

Transcription factors (TFs) are generally deemed undruggable due to their structural complexity. mRNA technologies have paved the way to overcome this therapeutic limitation by enabling the development of mRNA protein replacement therapies. Here we explore the newly described TF activity of LCOR (Ligand-dependent corepressor), which suppresses tumor growth by inducing the antigen presentation machinery (APM) of the tumor cells and constrains cellular plasticity. These LCOR effects facilitate recognition of the tumor by the immune system and immune-mediated tumor cell death. To deliver *Lcor* mRNA into tumor cells, we have used poly β-(amino esters) (pBAE) nanoparticles (NPs) for local delivery of *Lcor* mRNA in breast cancer primary tumor models. We have engineered pBAE-NPs with high potential for efficiently encapsulate mRNA and facilitate cellular uptake. Our results show optimal endosomal escape, which results in high transfection efficiency *in vitro* and *in vivo*, restoring LCOR function in tumor cells and engaging their APM. In preclinical triple-negative breast cancer (TNBC) models, the intratumoral delivery of *Lcor* mRNA led to a reduction in tumor growth. Importantly, the combination of *Lcor* mRNA-loaded NPs with anti-PDL1 or anti-CTLA4 immunotherapies eradicated most of the tumors in our preclinical TNBC model. Overall, our nanotherapeutic strategy emerges as an innovative TF-replacement therapy, leveraging the immunogenic effects of LCOR to eradicate breast cancer tumors when combined with immunotherapy.

**Gaphical Abstract:** 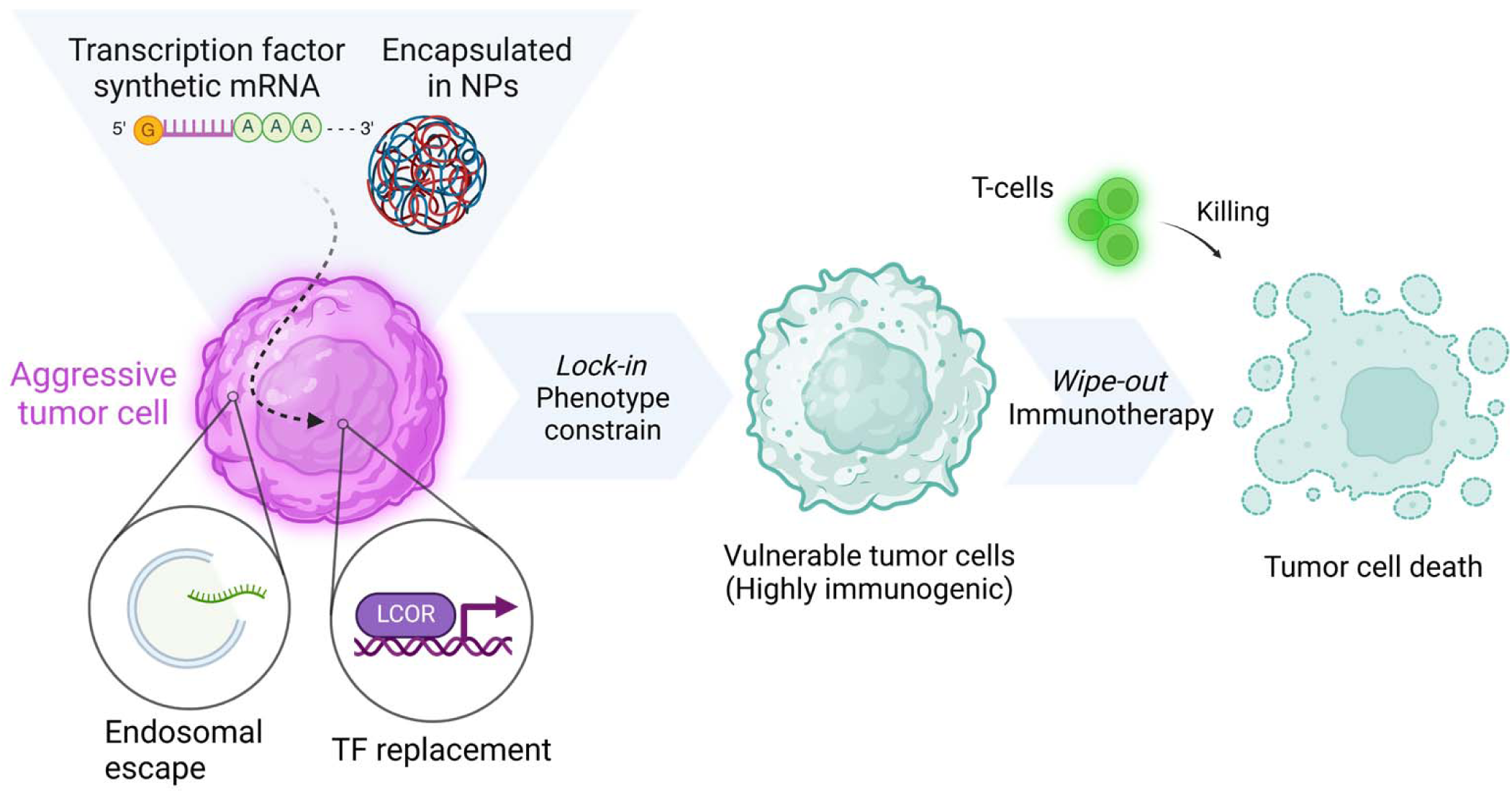

## Introduction

Transcription factors (TFs) have traditionally been considered undruggable, mostly because of their unstructured three-dimensional (3D) shape. This feature makes it almost impossible to target TFs with small molecules, making it challenging to modulate their activity for clinical applications^1^. In recent years, mRNA has emerged as a promising therapeutic biomolecule, bypassing the traditional drawbacks of small molecules and achieving unprecedented potential for a range of clinical needs, as evidenced by COVID-19 vaccines. mRNA technology is not only valuable for prophylactic vaccines but also for protein replacement therapies, which aim to substitute or restore specific protein in target cells, opening new avenues in modulating TFs activity to treat different disorders^2–5^. However, despite significant progress in mRNA technologies and nucleic acid delivery systems, achieving direct *in vivo* effective expression in target cells remains a major challenge in protein replacement therapies. Limitations mainly arise from mRNA stability in physiological fluids and low cell membrane penetration, which restrict success to the carrier and delivery system. Cationic polymer-based nanovectors are among the most promising candidates for mRNA deliver^6^. Polyplex formation relies on electrostatic interactions between the cationic amine groups of polymers and the polyanionic mRNAs. This interaction leads to polynucleotide condensation into nanosized particles with a positive surface charge, which protects the nucleic acid and enhances stability. Poly β-(amino esters) (pBAEs) polymers are a class of biocompatible and biodegradable polymers composed of amines and cleavable ester bonds in their backbones. These structures facilitate endosomal escape due to protonation of tertiary amines at lower pH^7^. The use of oligopeptide-end modified variants enhances their transfection capacity while preserving their non-toxic character^8,9^.

mRNA protein replacement approaches have been used in tumor cells to correct genetic alterations in preclinical models^2,3,8^. However, these approaches have not been used to modulate master TFs reprogramming cell phenotypes, such as differentiation, stemness, or immune evasion. We have previously described Ligand-dependent corepressor (LCOR) as a master TF regulating tumor cell immunogenicity in triple negative breast cancer (TNBC)^10,11^. LCOR increases the immune detection and immune mediated killing by orchestrating the transcriptional activation of the antigen processing and presentation machinery (APM) independently of interferon-gamma (IFN-γ) signaling^11^. This observation is relevant since the downregulation of the APM is a major cause of resistance to immune checkpoint blockade (ICB) therapy^12–14^, which is related to the loss of components of the IFN-γ signaling pathway^15–17^.

In breast cancer, ICB is part of the standard-of-care (SOC) for stage II/III TNBC and PDL1+ advanced TNBC^18,19^. Only anti-PD1/PDL1 therapy has been approved for TNBC, although others are under intense research, such as anti-CTLA4 therapy, which has shown moderately positive results in recent clinical trials in early and advanced TNBC^20,21^. However, in these patients, ICB leads to only a modest increase in response rates compared to chemotherapy alone^22^. Understanding and overcoming these barriers is critical for fully harnessing the potential of ICB therapies and improving outcomes for patients with TNBC.

Here we demonstrate proof-of-concept for the therapeutic modulation of breast cancer cell phenotypes using pBAE-nanoparticles (NPs) loaded with *Lcor* mRNA in preclinical mouse models. This approach showed exceptional efficacy in intratumoral settings, leading to tumor eradication when combined with PD-L1 or CTLA4 blockade therapies. These innovative therapeutic strategies open up new avenues for expanding the use of LCOR and TFs replacement therapies in immunotherapy and oncology.

## Methods

### Reagents

The reagents used for pBAE synthesis were bought from Sigma Aldrich® and the oligopeptides CH3 (NH2-Cys-His-His-His-COOH) and CR3 (NH2-Cys-Arg-Arg-Arg-COOH), with a purity higher than 95%, from Shanghai Yaxian Chemical Co. Ltd. All synthetic mRNAs were supplied by Cellerna, while the Cyanine3 and Cyanine5 NHS ester dyes (Cy3 and Cy5) were purchased from Lumiprobe.

### Synthesis of end-modified poly(β-aminoester) polymers

The pBAE C6 polymer used was synthesized by a Michael addition reaction, as previously described^23^. Briefly, 5-aminopentanol and 1-hexylamine were added to 1,4-butanediol diacrylate at the same ratio and using a slight excess of the latter (molar ratio of amine:dyacrilate, 1:1.1). The reaction was carried out at 90°C for 24 h. The polymer was further modified at its acrylate ends by the introduction of cationic oligopeptides, either CR_3_ or CH_3_ (molar ratio of polymer:peptide, 1:2.5). The mixture was stirred in dimethyl sulfoxide (DMSO) overnight at room temperature to later be precipitated in a mixture of diethyl ether/acetone. ^1^HNMR spectral characterization was performed to verify each synthesis step and the formation of the desired end structure.

### Nanoparticle formulation

The formulations were prepared using a mixture of polymer and nucleic acid, at a molar ratio of 25:1. Two starting solutions were prepared in sodium acetate buffer (12.5 mM, pH 5.2). One contained the mRNAs of choice at a final concentration of 0.5 mg/mL, while the other contained the polymer mixtures at a final concentration of 12.5 mg/mL. The polymer combinations used were C6-CR_3_ and C6-CH_3_ at a ratio of 60:40. The mRNA solution was pipetted into the polymer solution and thoroughly mixed for 15–20 sec, followed by a 30-min incubation at room temperature. Later, the solution was precipitated in DEPC water and a third of the final volume of HEPES buffer (20 mM, pH 7.4) with 4% (w/v) of sucrose was added^24^.

### mRNA labeling

The pH of the commercially available mRNA was first tested to determine whether it needed to be basified to achieve a neutral pH. The mRNA was mixed with Cy3 at a ratio of 10:3 (Cy3:mRNA) (V_1_) and stirred for 2 h at room temperature in the dark. Later, 0.1 times the V_1_ of a sodium chloride solution (5M) was added to the previous mixture (V_2_). Twice the V_2_ of cold pure ethanol was then added to the mixture and frozen overnight. The mixture was then centrifuged in a precooled centrifuge at 14,000x*g* at 4°C for 15 min. The supernatant was then removed, and the pellet was rinsed with 70% ethanol and centrifuged again several times until the supernatant was clear. The pellet was then left to dry and characterized using the nanodrop setting of an Infinite M Plex TECAN Plate reader, where the concentration and purity of the mRNA and the amount of Cy3 in the sample were calculated.

### Polymer labeling

The C6-CR_3_ at 100 mg/mL was mixed with a solution of Cy5 at 10 mg/mL in DMSO, taking 35 µL and 60 µL of each respectively. Triethylamine and DMSO were then added while protecting the reaction from light. The mixture was stirred for 20 h at room temperature. Next, it was precipitated in diethyl ether:acetone at a ratio of 7:3, vortexed, and centrifuged at 4000 rpm for 10 min. The supernatant was removed, and the pellet was rinsed twice with the same solution. The labeled polymer was left to dry for 24 h and then adjusted to 100 mg/mL in DMSO. Later, the amount of Cy5 was tested through a calibration curve with several dilutions of Cy5 and read by fluorescence in an Infinite M Plex TECAN Plate reader.

C6-CH3 labeling followed the same protocol but using 56 µL of C6-CH3 and 100 µL of Cy5.

### mRNA complexation capacity and stability

The complexation capacity of the mRNA in the NPs was tested using the Ribogreen kit. The NPs were formulated at a ratio of 60:40 (C6-CR_3_:C6-CH_3_) of the oligopeptide-modified polymers with EGFP mRNA as a reporter gene. For the test, fresh NPs (free mRNA) were prepared and diluted in buffer and also in buffer with heparin, causing NP disruption and thus total exposure of the encapsulated mRNA (total mRNA). The different NP conditions were then compared to three standard curves (mRNA range, heparin, and NPs with heparin). Following the equation below, the percentage of encapsulated mRNA or encapsulation efficiency was calculated.

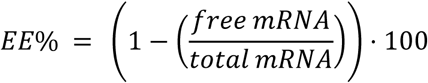

### Physicochemical characterization

NPs were characterized by Dynamic Light Scattering (DLS; ZetaSizer Nano ZS, Malvern). In this regard, their hydrodynamic diameter (nm) was measured, as well as their surface charge (mV) and poly-dispersity index (PDI), by taking measures at room temperature with a 633 nm laser wavelength and a detector at an angle of 173 °. The measurements were taken in triplicates, and the results are represented as mean ± standard deviation.

### Stability by Fluorescence Resonance Energy Transfer

NP stability was tested by fluorescence resonance energy transfer (FRET). To this end, the genetic material was labeled with Cy3 and the polymer with Cy5, thereby allowing the study of the complexation and structural organization of the NPs. Higher energy transfer indicates closer proximity between the fluorophores, thereby signifying the formation of a complexed NP. Conversely, a decrease in energy suggests the separation of the polymer and genetic material, leading to NP disassembly. FRET measurements were carried out using a TECAN Infinite M Plex microplate reader and a flat black 96-well plate. This test was performed in phosphate buffer saline (PBS) at 25°C, simulating laboratory stability, and in a solution of 50 mg/mL of Bovine Serum Albumin (BSA) at 37°C to simulate physiological stability.

### Cell culture

AT3, 4T07, 66cl4, 4T1, and EpRAS cells were kindly provided by Dr. Yibin Kang (Princeton University). All other cell lines were purchased from the American Type Culture Collection (ATCC). The identity of the cell lines used in this study was validated by short tandem repeat (STR) profiling and matched to reference databases to confirm authenticity. Dulbecco’s Modified Eagle’s Medium (DMEM; ATCC) was used to culture cells. The cell culture medium was supplemented with 10% fetal bovine serum (FBS), Pen-Strep (100 U ml−1), and L-Glutamine (100 μg ml−1). Experiments were performed at 37 °C in 5% CO_2_ conditions and the normal level of O_2_ in a cell culture incubator. All cell lines were routinely tested for Mycoplasma contamination.

### Cellular uptake of dye-labeled mRNA-loaded NPs

To monitor the cellular NP uptake, Cy3-*eGFP*-mRNA-Cy5-NPs and Cy3-ΔHTH-Lcor mutant-mRNA-Cy5-NPs were prepared. AT3 or 4T07 cells were first seeded in 96-well plates at a density of 1×10^4^ cells per well and incubated at 37 °C in 5% CO_2_ for 24 h. They were then incubated with medium (DMEM) containing Cy3-*eGFP*-mRNA-Cy5-NPs or Cy3-*Δhth*-mRNA-Cy5-NPs for different times. The cells were then washed with PBS, resuspended using Trypsin-0.25% EDTA

### Flow cytometry analysis

Flow cytometry data were collected on either a Fortessa Flow Cytometer (BD) using FACSDiva v.9.0 software and analyzed by FlowJo software v.10.8.1. Briefly, cells were trypsinized, washed twice with PBS, 5% FBS, and 0.5 mM EDTA, passed through a 70-μM filter, and counted before proceeding with a general staining protocol using anti-SIINFEKL (OVA) or anti-H-2K^d^.

### Immunoblotting

Protein extracts from cultured cells were lysed using RIPA buffer. Equal amounts of protein were determined using a bicinchoninic acid protein assay kit (Pierce/Thermo Scientific), following the manufacturer’s instructions. After gel electrophoresis and protein transference, membranes were blocked with 5% BSA in TBST for 1 h at room temperature with gentle shaking. Membranes were rinsed and then incubated overnight at 4 °C with appropriate primary antibodies. The immunoreactive bands were visualized using an enhanced chemiluminescence (ECL) detection system (Merck). Data was collected using Nine Alliance Q9 software in an UVITEC Cambridge system.

### RT–qPCR analysis

Total mRNA was purified using the RNeasy Mini Kit (Qiagen) and reverse transcribed into cDNA using the High-Capacity cDNA Reverse Transcription Kit (Life Technologies). RT–qPCR was performed using the LightCycler 480 SYBR Green I Master Mix (Roche). Data was collected using QuantStudio 12 K Flex software. mRNA expression was normalized by the expression of *Gapdh* (RT–qPCR primers are listed in Supplementary Table 1).

### Animals

Approximately 8-week-old female BALB/c and C57BL/6J mice were purchased from Charles River Laboratories and housed in a pathogen-free animal facility at the Barcelona Biomedical Research Park. All animal procedures presented in this study were approved by the Ethical Committee for Animal Research of the Barcelona Biomedical Research Park and the *Departament de Medi Ambient i Habitatge de la Generalitat de Catalunya* (Catalan Government). For all experimental procedures, euthanasia was applied once tumors reached 1,500 mm^3^ or when animal health was compromised. Five animals (with the same genotype and sex) were maintained in individually ventilated Makrolon cages at a temperature of 22 ± 2 °C, with a relative humidity of 55 ± 5%, 12 h light:dark cycle starting at 08:00 am, and food and water *ad libitum*.

### Orthotopic tumor model preparation

For primary tumor models, 5×10^3^ 4T07 cells or 5×10^5^ AT3 cells were orthotopically injected into the MFP using 1:1 PBS:Matrigel (Corning). Tumor volume was measured twice weekly with digital calipers, and calculations were applied (π × length × width^2^/6). In ICB assays, once tumors reached 30-50 mm^3^, they were treated with either pBAE-RH mRNA at a dose of 250 µg/kg by i.t. injection twice a week, anti-PDL1 at a dose of 5 mg/kg by i.p. injection once a week, or their combination. On the other hand, mice were treated with anti-CTLA4 with an initial dose of 5 mg/kg, followed by five doses at 0.5 mg/kg, by i.p. injection every 7 days, or their combination pBAE-RH mRNA at a dose of 250 µg/kg by i.t. injection twice a week.

### Tissue immunofluorescence

Paraffin-embedded tissue samples were cut into 3-μm slices. After tissue dehydration and deparaffinization, antigens were retrieved with 0.5 M citrate, pH 6, in a pressure bath for 20 min. Tissue slices were incubated in blocking buffer (PBS, 2.5% Normal Goat Serum, 0.1% Triton-X) and incubated overnight at 4°C with primary antibody. Tissue was washed and fluorescent conjugated secondary antibodies were incubated for 1 h at room temperature. It was then washed again and samples were mounted in DAPI medium (0100-20, SouthernBiotech).

### Fluorescent in situ hybridization (FISH)

Paraffin-embedded tissue samples were cut into 3-μm slices. After tissue dehydration and deparaffinization, samples were treated for 10 min with hydrogen peroxide. Antigens were retrieved with 0.5 M citrate (pH 6) in a pressure bath for 20 min. Tissue slices were treated for 30 min with Protease solution at 40°C. The RNAscope^TM^ 2.5 HD detection kit was then used following the manufacturer’s instructions (322360, ACD). *Lcor* mRNA was detected using mouse Lcor RNAscope^TM^. Samples were washed and mounted in DAPI medium.

### pBAE-NP transfection efficacy *in vivo*

Once tumors reached the desired volume, mice were euthanized and tumors were extracted. Samples were mechanically digested followed by a 30-min enzymatic digestion with 2 mg/ml Collagenase A and 0.5 mg/ml DNAse I. Cells were passed through a 100-μm strainer and incubated for 5 min at 37°C in ACK lysing buffer. They were then passed through a 40-μm strainer, counted, and processed for flow cytometry analysis.

### Statistical analysis

The sample size of the animal studies was determined based on pilot experiments or previous studies. Experiments were performed with female mice of a similar age. Mice were randomized before cell injections and treatment allocation. Researchers were not blinded during experimentation and outcome since it was necessary to monitor each group. For all *in vivo* and *in vitro* experiments, independent biological replicates are indicated in the figure legends. Results are represented as mean ± S.E.M. Normality of data was checked using the Shapiro test and homoscedasticity using Barlett’s test. For two groups of parametric data, statistical significance was assessed using the Mann-Whitney U test. For multiple independent groups, one way ANOVA, the Kruskall-Wallis H test or Multiple Mixed-effect analysis was used. For statistical significance. *p<0.05, **p<0.01, ***p<0.001. All experiments were reproduced in at least three independent biological replicates unless specified in the figure legend.

## Results

### mRNA delivery by pBAE-NPs *in vitro*

pBAE-NPs were formed by the electrostatic interaction of mRNA with the polymer, using a polymer combination of 60% arginine (R) and 40% histidine (H) oligopeptides as cationic end-modified pBAEs (Fig. 1A), previously confirmed to be non-toxic^25^. NP synthesis yielded small nanometric and monodisperse particles (Fig. 1B and Fig. S1A) with high transfection efficiency *in vitro* and *in vivo* in several hosts ^8,26^. The efficiency of *eGFP* mRNA-loaded pBAE-NPs to transfect mRNA into different mouse breast cancer cells (AT3, 4T07, EO771, EMT6, 66cl4, EpRAS, and 4T1) was tested using NPs encapsulating *eGFP* mRNA, at a dose of 2.5 µg/mL. After 48h treatment, AT3 and 4T07 cells displayed the highest translated <u>eGFP</u> protein after NP uptake (Fig. S1B). In human models MDA-

**Figure 1.**
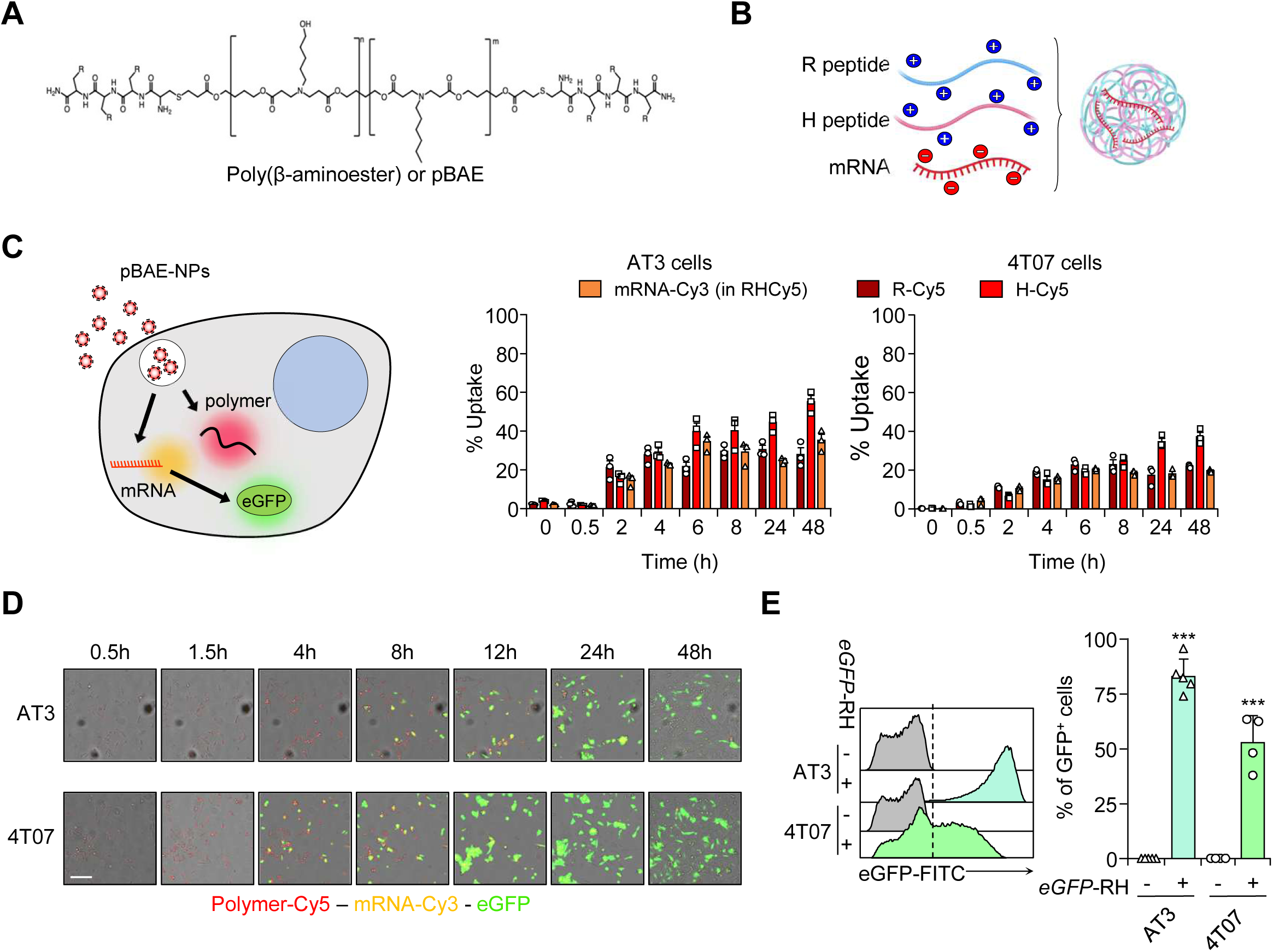
pBAE-NPs efficiently transfect breast cancer cell lines with mRNA *in vitro*. **(a)** Poly β-(amino ester) (pBAE) polymers are composed of cleavable ester bonds and amines. The R terminal can be an arginine or histidine oligopeptide. **(b)** Polyplex assembly is based on interactions between the positively charged amine groups of polymers and the negatively charged mRNA. This interaction leads to condensation into nanosized particles, which protect the nucleic acid. **(c)** Flow cytometry analysis of the percentage of pBAE-NP uptake over time in TNBC 4T07 and AT3 mouse cell lines. mRNA was labeled with cyanine (Cy)-3, whereas the polymer was labeled with Cy5. Data represent the mean % of total gated cells + S.E.M. N = 3 independent experiments. **(d)** Fluorescence microscopy images of TNBC 4T07 and AT3 cells treated with pBAE-NPs labeled with Cy3 (eGFP-mRNA) and Cy5 (R-modified polymer) over time. Scale bar = 50 µm. **(e)** Flow cytometry analysis of the percentage of TNBC 4T07 and AT3 cells positive for eGFP expression from mRNA-loaded pBAE-NPs. Data represent the mean % of total gated cells + S.E.M. N = 4–5 independent experiments. Statistical significance; *p<0.05, **p<0.01, ***p<0.001, by one-way ANOVA test in C and Kruskal-Wallis H test followed by Dunn’s post hoc test in E.

MB-231 and MCF7 cells, the NPs also showed high *eGFP* mRNA transfection efficiency (Fig. S1C). pBAE-NPs efficiently encapsulated 60% of *Lcor* mRNA (Fig. S1D). To determine the endosomal escape of these NPs across the different cell lines, we fluorescently labeled the polymer and mRNA with Cyanine (Cy)5 and Cy3, respectively, and monitored endosomal escape and mRNA translation kinetics by flow cytometry and fluorescence microscopy. Time-lapse imaging analysis of Cy5 and Cy3, and also eGFP intensity revealed the endosomal escape and translation rate between cell lines, the 4T07 cells showing higher overall mRNA delivery efficiency than AT3 cells (Fig. 1C, D). Flow cytometry analysis of eGFP fluorescence at 12 h showed higher uptake and translation of *eGPF* mRNA in AT3 cells (Fig. 1E), with the expected dose-dependent effect (Fig. S1E). These results demonstrate that tumor cells can uptake the mRNA-loaded pBAE-NPs and that these particles can escape the endosomal system and deliver the mRNA to the cytoplasm for proper translation of the protein.

### *Lcor* mRNA delivery by pBAE-NPs and TF functionality in tumor cells *in vitro*

To test the dynamics and ability of our system to deliver TFs to tumor cells, we treated AT3 and 4T07 cells *in vitro* with pBAE-NPs loaded with *Lcor* mRNA. Synthetic *Lcor* mRNA contained a Cap1, 5’ and 3’ untranslated regions (UTR) and a standard polyA tail (Fig. S2A), and all uracil were replaced for 5-methoxyuracil (5-moU) to avoid immunogenic reactions^27,28^. First, we measured and detected high levels of *Lcor* mRNA by qRT-PCR (Fig. 2A, B), increasing in a dose-dependent manner (Fig. S2B), as expected. pBAE NPs were stable at 25°C for 24 h (Fig. S2C). In contrast, under conditions simulating the physiological environment (37°C), a decrease in FRET signaling was detected as the incubation time increased, indicating disassembly of the NPs after 2 h (Fig. S2C). After treatment of the cells with the NPs *in vitro*, we detected cellular uptake of mRNA, with a peak at 6 h, which was maintained up to 36 h in culture (Fig. 2A, B), thereby indicating a rapid uptake and release of mRNA and a transient cell expression. Importantly, to measure the translated product of the *Lcor* mRNA delivered, we performed western blot analysis. High levels of LCOR protein were observed in tumor cells after 12 h of RH pBAE-NPs-*Lcor* mRNA treatment (Fig. 2C). These findings indicate that this technology can deliver synthetic *Lcor* mRNA into tumor cells, where it is efficiently translated into LCOR protein.

**Figure 2.**
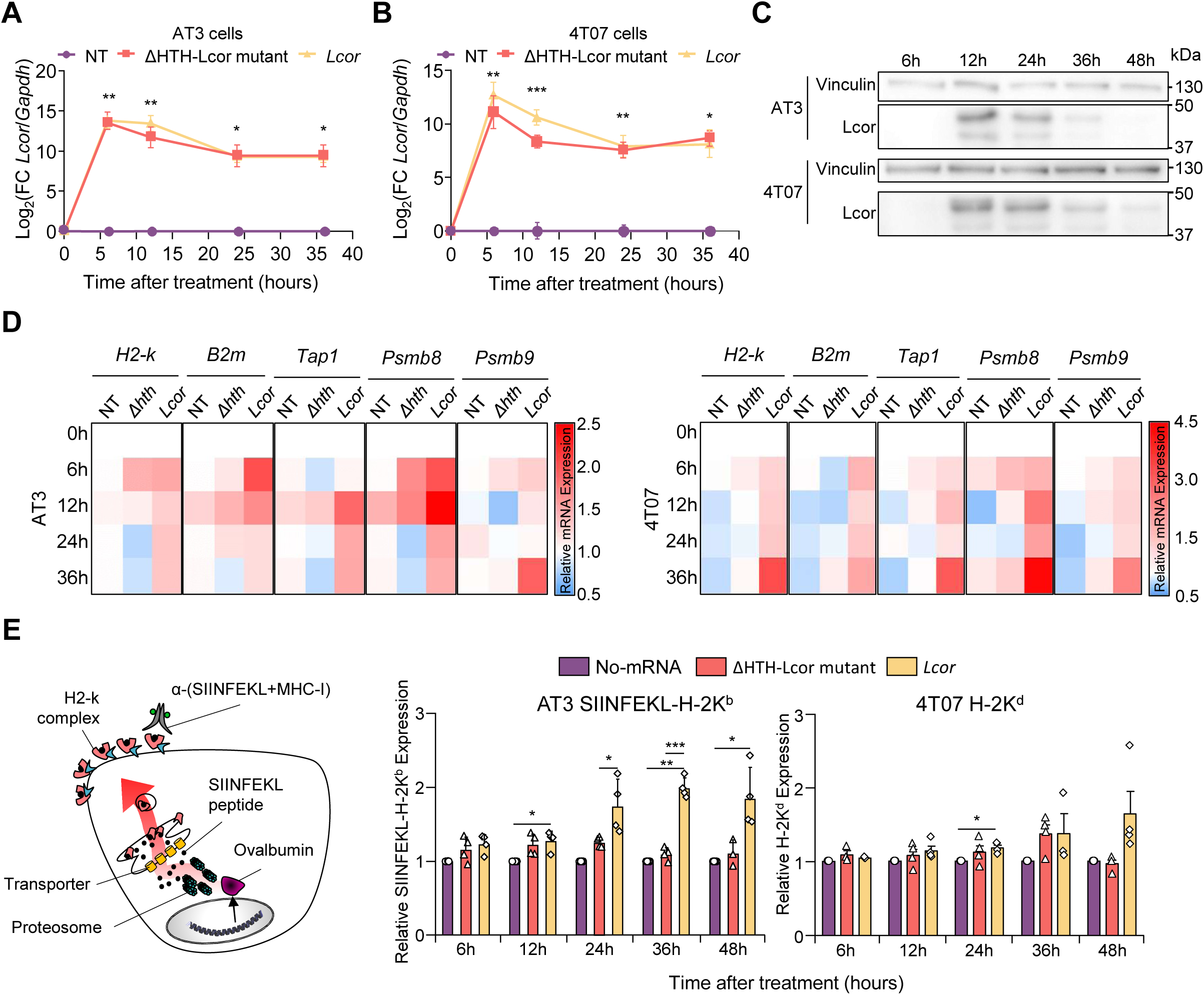
*Lcor* mRNA delivery by pBAE-NPs and TF functionality in tumor cells *in vitro*. **(a)** RT-qPCR analysis of *Lcor* mRNA levels over time in TNBC AT3 cells treated with the indicated mRNA. Data are represented as Log_2_(Fc) to the normalized expression of non-treated control condition. Expression levels were normalized to *Gapdh.* N = 4 independent experiments. Data represent the mean ± S.E.M. **(b)** RT-qPCR analysis of *Lcor* mRNA levels over time in TNBC 4T07 cells treated with the indicated mRNA. Data are represented as Log_2_(Fc) to the normalized expression of non-treated control condition. Expression levels were normalized to *Gapdh*. N = 3 independent experiments. Data represent the mean ± S.E.M. **(c)** Western blot image of TNBC AT3 and 4T07 cells treated with *Lcor* mRNA-loaded pBAE-NPs for 6, 12, 24, 36, and 48 h. Detection of Lcor and Vinculin (loading control). **(d)** RT-qPCR analysis of APM genes over time in TNBC AT3 (left) and 4T07 (right) cells treated with the indicated mRNA. Data are represented as Fc to the normalized expression of non-treated control condition. Expression levels were normalized to *Gapdh*. N = 4 independent experiments for AT3 and 3 independent experiments for 4T07 Data represent the mean. **(e)** Flow cytometry analysis of antigen presentation levels in AT3-OVA and 4T07 cells treated with the indicated mRNAs for 6, 12, 24, 36, and 48 h. AT3-OVA cells were stained for SIINFEKL-H-2K^b^ complexes and 4T07 cells for H-2K^d^ to assess peptide-MHC I presentation. N = 4 independent experiments. Statistical significance; *p<0.05, **p<0.01, ***p<0.001, by linear mixed model analysis with Geisser-Greenhouse correction, followed by Tukey’s multiple comparisons test.

Next, to assess whether LCOR can restore its function as an inducer of APM activity in tumor cells, we evaluated APM gene expression at different time points *in vitro* in AT3 and 4T07 cells treated with a single dose of 5 μg of *Lcor* mRNA-loaded RH pBAE NPs. The ΔHTH-Lcor mutant mRNA was used as negative control (Fig. S2A). This mutant lacks the HTH DNA binding domain, which is responsible for directing the protein to induce APM gene expression^11^. The functional results showed that *Lcor* mRNA NPs, induces the expression of APM genes in AT3 and 4T07 cell lines. The results showed a functional effect of the NP treatment mediated by LCOR-inducing APM genes in AT3 and 4T07 cell lines (Fig. 2D and S2D), thus reconfiguring the immunogenic phenotype of the cells. Next, we used AT3 cells that constitutively overexpress ovalbumin (OVA). In these cells, OVA is cleaved, generating the SIINFEKL antigen peptide presented in the H-2K^b^ context. This can be used to measure APM activity using the anti-SIINFEKL antibody via flow cytometry^29^. The results showed an increased number of cells with higher OVA-SIINFEKL presentation, indicating the enhanced activity of the APM induced by the *Lcor* mRNA-loaded pBAE-NPs. We also observed a time- and dose-dependent effect regarding APM induction (Fig. 2E and Fig. S2E). Overall, these results demonstrate the efficiency of this mRNA nanotechnology to rescue the function of the LCOR TF in inducing tumor cell immunogenicity and thus modulating tumor phenotypes^11^.

### pBAE-NPs efficiently deliver mRNA into breast cancer tumors *in vivo*

To test the capacity of pBAE-NPs to deliver mRNA to tumor tissue from preclinical models, we encapsulated *eGFP* or *Firefly Luciferase* (*FLuc*) mRNA for fluorescence or bioluminescence (BLI) detection. First, we generated primary tumors by orthotopic mammary fat pad (MFP) injection of AT3 or 4T07 cells into immunocompetent C57BL/6 or BALB/c mice, respectively. When tumors reached 0.5 x 0.5 cm^2^, we treated them intratumorally with pBAE-NPs loaded with 5 ug of synthetic *FLuc* or *eGFP* mRNA. We detected BLI at 3 h, meaning that tumor cells had taken up the mRNA-loaded NPs and translated a luciferase active protein within 3 h. In both models, expression peaked around 6 to 10 hours after administration (Fig. S3A, B). In addition, we injected *eGFP* mRNA-loaded pBAE-NPs into mice bearing 4T07-mCherry tumors. After 6 h, we harvested and digested the tumor tissues to evaluate, by GFP fluorescence using flow cytometry, the number of tumor or stromal cells that had efficiently taken up the NPs and translated their mRNA cargo. Both tumor cells (40–80%) and stromal cells (30–60%) incorporated and translated the mRNA (Fig. 3A), thereby indicating substantial transfection capacity *in vivo* after one dose of 5 μg mRNA-loaded NPs.

**Figure 3.**
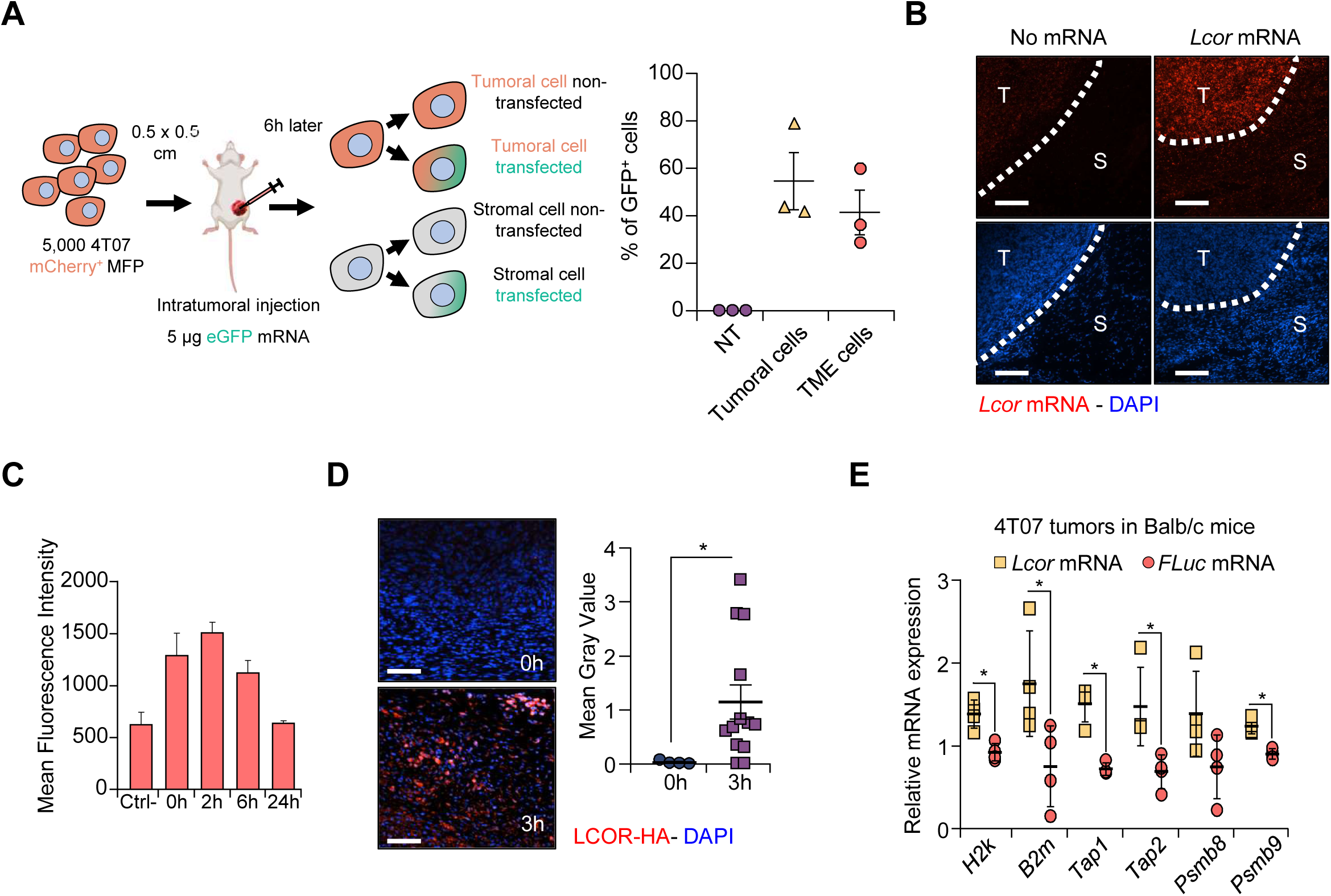
pBAE-NPs efficiently deliver mRNA into breast tumors *in vivo*. **(a)** Flow cytometry analysis of eGFP mRNA uptake in tumor (mCherry⁺) and stromal (mCherry⁻) cells. Once tumors reached 0.5 × 0.5 cm^2^, tumors were intratumorally treated with 5 µg of eGFP mRNA-loaded pBAE-NPs. After 6 h, the tumors were harvested, mechanically and enzymatically digested for single-cell suspension, and analyzed. Data represent the % of eGFP⁺ cells in each population. Results are shown as mean ± S.E.M. from N = 3 mice. **(b)** Representative fluorescence images of *Lcor* mRNA detected by in situ hybridization (ISH) (left) and nuclei stained with DAPI (blue). After reaching a size of 0.5 × 0.5 cm^2^, tumors formed by 4T07 cells implanted in the mammary fat pad of BALB/c tumors were intratumorally treated with 5 µg of *Lcor* mRNA-loaded pBAE-NPs. Quantification of mean fluorescence intensity (MFI) of *Lcor* mRNA at 0, 3, 6, and 24 h post-treatment (right). Data are shown as mean ± S.E.M. from N = 3 mice per time point. Scale bar = 40 µm **(c)** Representative fluorescence images of Lcor-HA protein detected with anti-HA antibody 3 h post-treatment. Nuclei are stained with DAPI (blue). After reaching a size of 0.5 × 0.5 cm^2^, tumors formed by 4T07 cells implanted in the mammary fat pad of BALB/c mice were intratumorally treated with 5 µg of *Lcor* mRNA-loaded pBAE-NPs. Quantification of fluorescence intensity was determined by ImageJ, showing the mean gray value. Data are shown as mean ± S.E.M. Images are representative of N = 3 mice. Scale bar = 30 µm **(d)** RT-qPCR analysis of APM genes in tumors formed by 4T07 cells implanted in the mammary fat pad of BALB/c mice. After reaching a size of 0.5 × 0.5 cm^2^, tumors were intratumorally treated with 5 µg of *Lcor* mRNA-loaded pBAE-NPs and then harvested 24 h post-treatment. Data represent the Fc of APM genes, normalized to *Gapdh*, shown as mean ± S.E.M. from N = 5 mice per group. Statistical significance: *p<0.05, **p<0.01, ***p<0.001, by Kruskal-Wallis H test followed by Dunn’s post hoc test in B, and Mann-Whitney U test in C and D.

Next, we tested the detection and kinetics of *Lcor* mRNA delivery *in vivo*. To this end, we used specific *Lcor* probes for tissue *in situ* hybridization (ISH) and immunofluorescence (IF) (Fig. 3B). After local administration of 5 μg of *Lcor* mRNA-loaded NPs, we observed a rapid increase in *Lcor* mRNA in the tumor tissue, followed by a decrease, reaching baseline levels after 24 h (Fig. 3C). Consistently, similar results were obtained by RT-qPCR (Fig. S3C, D). To unravel the protein dynamics, we used synthetic *Lcor* mRNA followed by the HA tag sequence to generate LCOR-HA protein and uniquely detect the ectopic protein using anti-HA by IF. As expected, LCOR-HA protein expression was delayed, peaking 3 h after administration (Fig. 3D). Linked to protein expression, at 3 h and 6 h after administration, we detected an increase in APM genes by RT-qPCR (Fig. 3E and S3D). Based on these results, we estimated an optimal therapeutic regimen of *Lcor*-mRNA-loaded pBAE-NPs administration in our preclinical experimental models would be every 3 days.

### *Lcor* mRNA nanotherapy synergizes with immune checkpoint inhibitors in preclinical models

To test the efficacy of the TF-replacement therapy using *Lcor*-mRNA-loaded pBAE-NPs, we first generated 4T07 breast primary tumors in immunocompetent BALB/c mice. Once tumors reached 0.4 x 0.4 cm^2^, *Lcor* mRNA-loaded NPs were administered at a dose of 250 µg/kg by intratumoral (i.t.) injection twice a week. In addition, we combined the *Lcor* mRNA therapy with the anti-PDL1 immune-checkpoint inhibitor (ICI), at a dose of 5 mg/kg, by intraperitoneal (i.p.) injection once a week (Fig. 4A). The combined *Lcor* mRNA-loaded NPs plus anti-PDL1 therapy did not have additional adverse immune effects, and mouse weights were similar across conditions during the treatment (Fig. S4A). Importantly, the results revealed that *Lcor* mRNA monotherapy was enough to reduce 4T07 tumor growth (Fig. 4A, B, and Fig. S4B). Furthermore, the combination of *Lcor* mRNA-loaded NPs with anti-PDL1 therapy not only reduced tumor growth but also led to tumor eradication in 5 out of 7 mice (Fig. 4B, D), and significantly increased the overall survival of treated animals (Fig. 4C). These results indicate that our *Lcor* mRNA nanotherapy synergized with anti-PDL1 and overcame resistance in 4T07 tumors.

**Figure 4.**
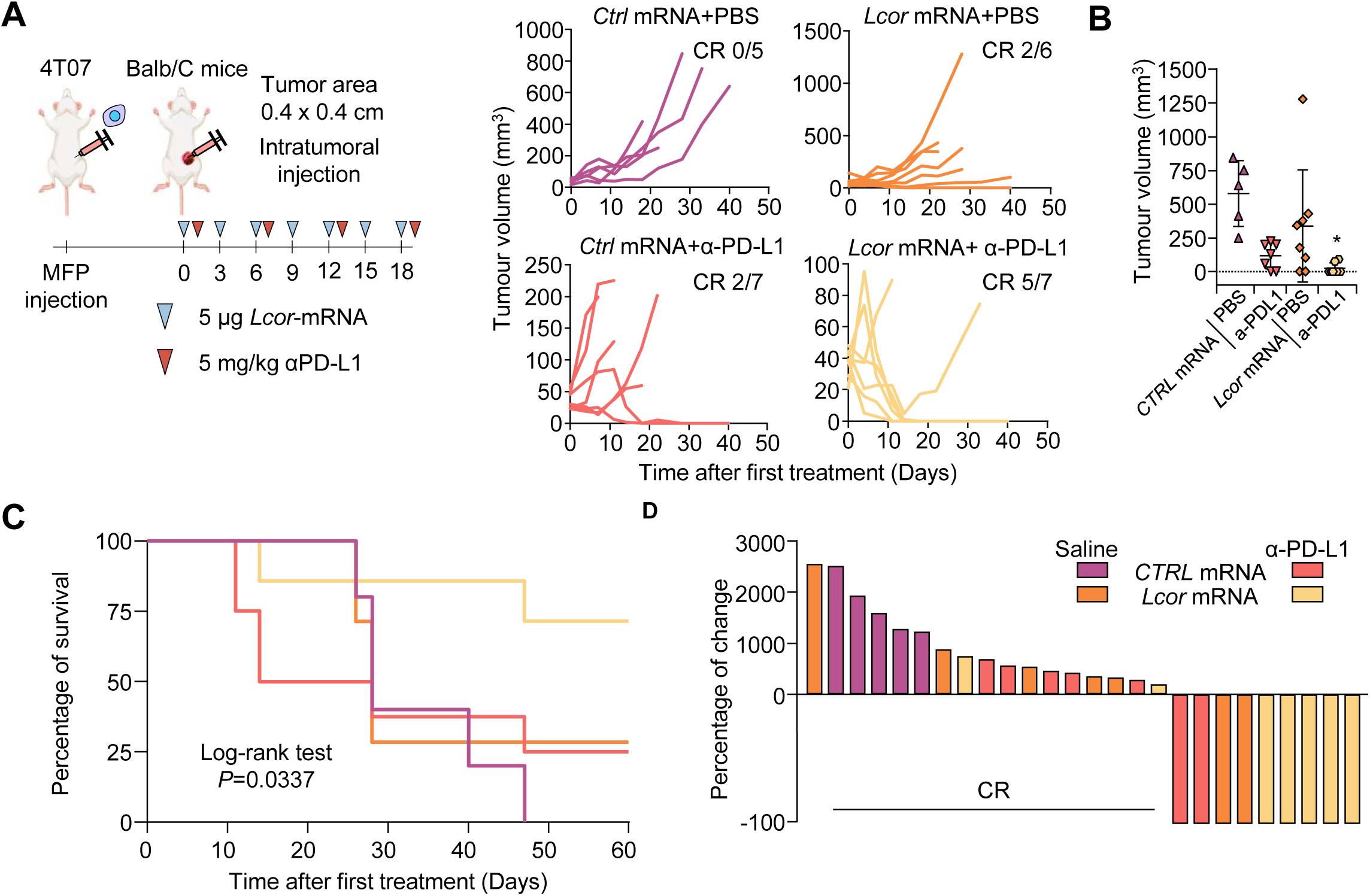
*Lcor* mRNA nanotherapy synergizes with anti-PD-L1 in preclinical models. **(a)** Tumor volume over time following the specified treatments. Mice were implanted with 4T07 cells in the mammary fat pad and treated with 5 µg of *Lcor* mRNA-loaded pBAE-NPintratumorally twice a week, and 5 mg/kg of anti-PD-L1 intraperitoneally once a week. **(b)** Tumor volume at the endpoint of the experiment for each treatment group. Data represent mean ± S.E.M. from N = 5 to 7 mice per group. Statistical significance: *p<0.05, **p<0.01, ***p<0.001, by Kruskal-Wallis H test, followed by Dunn’s post hoc test. **(c)** Kaplan-Meier survival curves with statistical analysis by log-rank test. **(d)** Waterfall plot of the percentage change in tumor volume for each mouse.

We extended our therapeutic approach to combine our *Lcor* mRNA NP therapy with the ICI anti-CTLA4. Mice were subjected to MFP implantation of 4T07 cells. When primary tumors reached 0.4 x 0.4 cm^2^, the animals received either *Lcor* mRNA-loaded NPs at a dose of 250 µg/kg by i.t. injection twice a week alone or in combination with anti-CTLA4, at an initial dose of 5 mg/kg, followed by five doses at 0.5 mg/kg by i.p. injection every 7 days (Fig. 5A). All the mice treated with the combination therapy showed a complete response (Fig. 5A, C and Fig. S5A), and thus increased survival (Fig. 5B).

**Figure 5.**
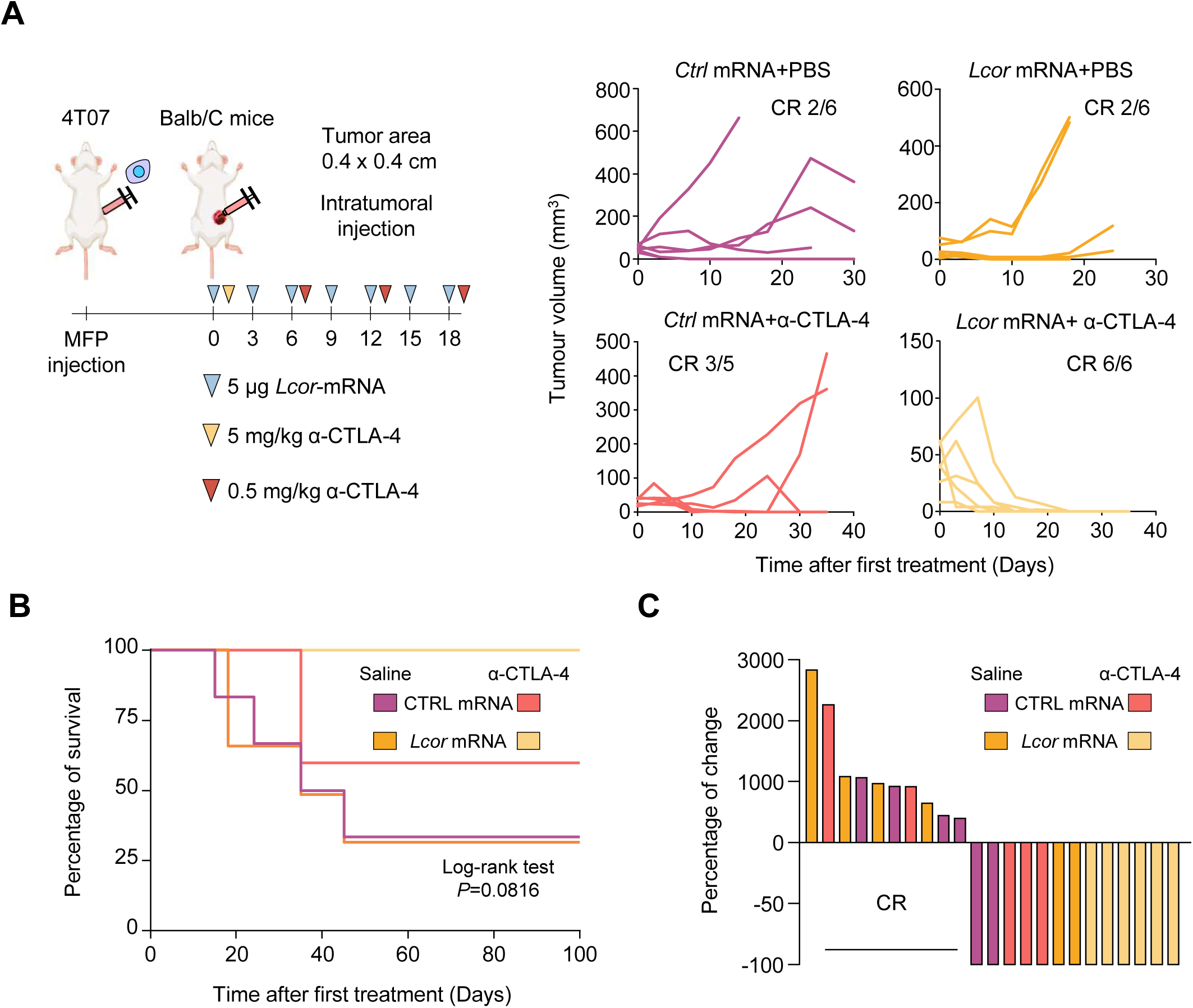
*Lcor* mRNA nanotherapy synergizes with anti-CTLA4 in preclinical models. **(a)** Tumor volume over time following the specified treatments. Mice were implanted with 4T07 cells in the mammary fat pad and treated with 5 µg of *Lcor* mRNA-loaded pBAE-NP intratumorally twice a week, and 5 mg/kg of anti-CTLA4 intraperitoneally for the first dose, followed by 0.5 mg/kg for subsequent doses. **(b)** Kaplan-Meier survival curves with statistical analysis by log-rank test. **(c)** Waterfall plot of the percentage change in tumor volume for each mouse. Data are shown from N = 5–6 mice per group, with results from two independent experiments.

Overall, these results demonstrate the potential of our strategy in leveraging mRNA therapy to rescue the LCOR TF. The combination of *Lcor* mRNA-loaded NPs with different ICIs showed high efficiency in preclinical models, thus supporting the feasibility of starting clinical studies and thus bringing the treatment closer to patients.

## Discussion

Here we provide evidence of the capacity of nanomedicine technologies to target TFs and reprogram cellular phenotypes into immunogenic tumor cell phenotypes. The identification of phenotype drivers or master TFs establishing core regulatory circuits and molecular memory presents potential opportunities for cell reconfiguration^30,31^. However, master TFs are usually non-druggable^1^. We overcame this therapeutic limitation by leveraging emergent RNA technologies and pBAE-NPs, in combination with immunotherapy, to enhance LCOR expression in tumors and thus promote tumor eradication. These observations lead to a paradigm shift for driver-cell state or driver-phenotype therapies in cancer.

mRNA therapies emerge as versatile tools with the potential to transform medicine. Among the different types of mRNA therapy, which are focused mainly on personalized neoantigen vaccines, tumor suppressor replacement therapies of mutated p53 and PTEN have been used in preclinical studies^32,33^. In our case, we modified LCOR expression, which is not mutated but downregulated in the most aggressive tumor cell populations, including immune-evasive cancer stem cells^11^. In this regard, we are the first to fine-tune tumor cell phenotypes with mRNA therapy based on a TF. This new approach of harnessing mRNA technologies for cancer therapy holds promise in oncology and immuno-oncology. Moreover, a potential transversal impact on other disciplines of medicine, such as immune-related disease and regenerative medicine, is also expected.

To achieve the functionality of exogenously administered mRNAs, an advanced delivery system is required. Among the different approaches available, polymeric NPs—specifically pBAE polyplexes— show a high transfection efficiency for the *in vivo* expression of diverse encapsulated nucleic acids with low toxicity^34–38^. In our preclinical models, these pBAE-NPs effectively transfected a range of breast cancer cells both *in vitro* and *in vivo*, demonstrating their potential as versatile delivery vehicles. pBAE-NPs offer advantages, including enhanced biocompatibility and biodegradability, chemical versatility, improved endosomal escape, reduced immunogenicity, effective targeting of extrahepatic tissues, suitability for local delivery, and ease of manufacture with greater stability ^6,39^. We observed cell-type dependence of NP uptake across the breast cancer lines tested, which could lead to differential efficiency in distinct types of tumors. Cell-dependence uptake was previously reported by our group and attributed to the differential endocytic pathways used by specific cell types^25,26^. In addition, pBAE-NPs are more prone to disassemble under physiological conditions than other NPs, i.e., they efficiently release the mRNA in response to a decrease in pH within the endosomal pathway, thus facilitating endosomal escape. This avoids eventual toxicity caused by NP accumulation, a phenomenon observed in other types of nanomaterial. However, NP disassembly could also compromise the efficacy of systemic administration. Our study shows that local administration reduces systemic toxicity, and the effective dose needed, while also lowering manufacturing costs since specific targeting is not required. Thus, this delivery mode is suitable for local neoadjuvant or intratumoral local therapies, similar to the use of NPs in protein replacement therapies^40^.

Our results highlight not only the potential of LCOR nanotherapy as a standalone treatment but also when combined with ICB therapies. Given that LCOR strongly induces tumor cell immunogenicity and immune recognition, our nanotherapy is designed to efficiently enhance immunogenicity. As such, LCOR is an ideal complement to ICB agents designed to restore T cells^11^. In fact, in immuno-oncology, various combination therapies, including chemotherapy, are used as induction treatments to increase immunogenicity, causing DNA damage and genomic instability^41,42^. Here we propose *Lcor* mRNA therapy to specifically instigate immunogenic functions that could replace the need to use chemotherapeutic induction treatment strategies. Another breakthrough of our study is that we demonstrate, for the first time, the remarkable performance of *Lcor* mRNA therapy combined with anti-CTLA4, as well as preclinical benefit of using *Lcor* mRNA-loaded NPs combined with anti-PD-L1 blockade. Of note, anti-CTLA4-based therapies have received less attention than anti-PD-L1 in breast cancer and none has been approved for clinical practice to date. However, new clinical trials have demonstrated the preliminary efficacy of anti-CTLA4 blockade in early-TNBC without the administration of chemotherapy^21^. Therefore, our results indicate the potential benefit of combining our nanotherapy with anti-CTLA4 blockade, without the need for chemotherapy.

Overall, the therapeutic approach described herein shows great potential to modulate the TFs that govern cancer cell fates. We envision multiple applications adapted from this technology to regulate malignant cell states determined by dysregulated TFs, thus providing new opportunities to tackle the emerging paradigm of cancer: driver phenotypes. Our study exemplifies this potential. Our future research will explore the suitability of *Lcor* mRNA therapy for other types of cancer, particularly poorly immunogenic and low MHC tumors, with the goal of reconfiguring and enhancing their immunogenicity to ensure immunotherapeutic clinical benefits.

## List of abbreviations

1HNMR: Proton nuclear magnetic resonance
3D: Three-dimensional
5-moU: 5-methoxyuracil
APM: Antigen presentation machinery
ATCC: American Type Culture Collection
BSA: Bovine Serum Albumin
CTLA4: Cytotoxic T-lymphocyte associated protein 4
Cy: Cyanine
DBD: DNA binding domain
DLS: Dynamic light scattering
DMEM: Dulbecco’s Modified Eagle’s Medium
DMSO: Dimethyl sulfoxide
ECL: Enhanced chemiluminescence
eGFP: Enhanced green fluorescent protein
FBS: Fetal bovine serum
Fluc: Firefly luciferase
FRET: Fluorescence resonance energy transfer
HTH: Helix-turn-helix
i.p.: Intraperitoneal
i.t.: Intratumoral
ICB: Immune checkpoint blockade
ICI: Immune-checkpoint inhibitor
IFN-γ: Interferon-gamma
LCOR: Ligand-dependent corepressor
NP: Nanoparticle
OVA: Ovalbumin
pBAE: Poly β-(amino esters)
PBS: Phosphate buffer saline
PDI: Poly-dispersity index
PDL1: Programmed Death-ligand 1
SOC: Standard-of-care
STR: Short tandem repeat
TF: Transcription factor
TNBC: Triple negative breast cancer
UTR: Untranslated regions

## Declarations

### Ethics approval

All animal procedures presented in this study were approved by the Ethical Committee for Animal Research of the Barcelona Biomedical Research Park and the *Departament de Medi Ambient i Habitatge de la Generalitat de Catalunya* (Catalan Government).

### Consent for publication

Not applicable

### Availability of data and materials

In this study no public datasets or proprietary datasets were used or generated.

### Author contributions

T.C.-T. devised the concept of the study and supervised the project with C.F. T.C.-T, C.F., and G.S.-M designed the study. G.S.-M. performed most of the *in vitro* and all the *in vivo* experiments, with the help of J.A.P, S.B.-B., A.L.-P., and M.S.-F. E.H.-M. and M.S. synthesized and biophysically characterized all the NPs in this study and performed some of the *in vitro* experiments. C.C. performed the microscopy imaging. All authors reviewed the manuscript.

## Acknowledgments

The authors thank C.R. for valuable discussions on the results and A.C.M. and M.N.L. for their technical support during the course of this study. Financial support from the Institute of Health Carlos III (ISCIII) (AC22/00042) and FCAECC (Award no. TRNSC213882FORN) within the TRANSCAN framework and by the Joint Transnational Initiative ERA-NET TRANSCAN-3, the European Commission; from MICIN/AEI and FEDER/UE (Proyecto PID2021-125910OB-I00, funded by /10.13039/501100011033); and from Generalitat de Catalunya (2021 SGR 00537) is acknowledged.

## Supplementary FIGURE LEGENDS

**Figure S1.**
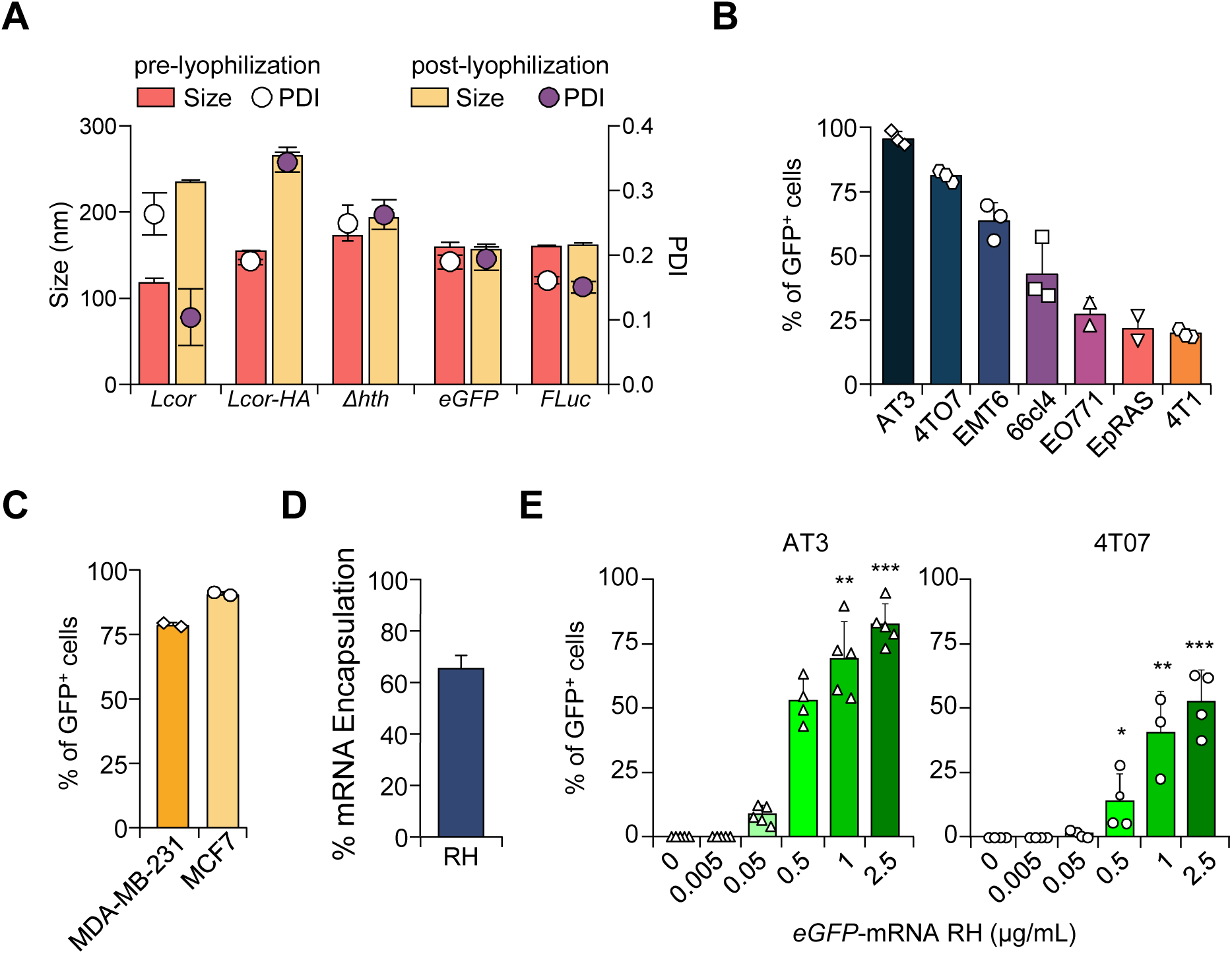
**(a)** Dynamic light scattering (DLS) analysis of nanoparticles (NPs) was used in this study. Size and polydispersity index (PDI) were measured before and after lyophilization. Data represent mean ± S.E.M. from 3 independent measurements. **(b)** Flow cytometry analysis of the percentage of several TNBC cells positive for eGFP expression after treatment with mRNA encapsulated in pBAE-NPs. Cells were treated with 5 µg of mRNA-loaded pBAE-NPs for 48 h. Data represent mean % of total gated cells + S.E.M. N = 2–3 independent experiments. **(c)** Flow cytometry analysis of the percentage of human TNBC MDA-MB-231 and ER+BC MCF7 cells positive for eGFP expression from mRNA-loaded pBAE-NPs. Cells were treated with 5 µg of mRNA-loaded pBAE-NPs for 48 h. Data represent mean % of total gated cells + S.E.M. N = 2 independent experiments. **(d)** Percentage of *Lcor* mRNA encapsulated by pBAE-NPs measured by fluorescence after treatment with the commercial Quant-it^TM^ RiboGreen kit. Data represent mean + S.E.M. **(e)** Flow cytometry analysis of the percentage of TNBC 4T07 and AT3 positive for eGFP expression after treatment with mRNA-loaded pBAE-NPs. Cells were treated with the dose indicated of mRNA-loaded pBAE-NPs for 12 h. Data represent mean % of total gated cells + S.E.M. N = 3–5 independent experiments. Statistical significance: *p<0.05, **p<0.01, ***p<0.001, by Kruskal-Wallis H test, followed by Dunn’s post hoc test in E.

**Figure S2.**
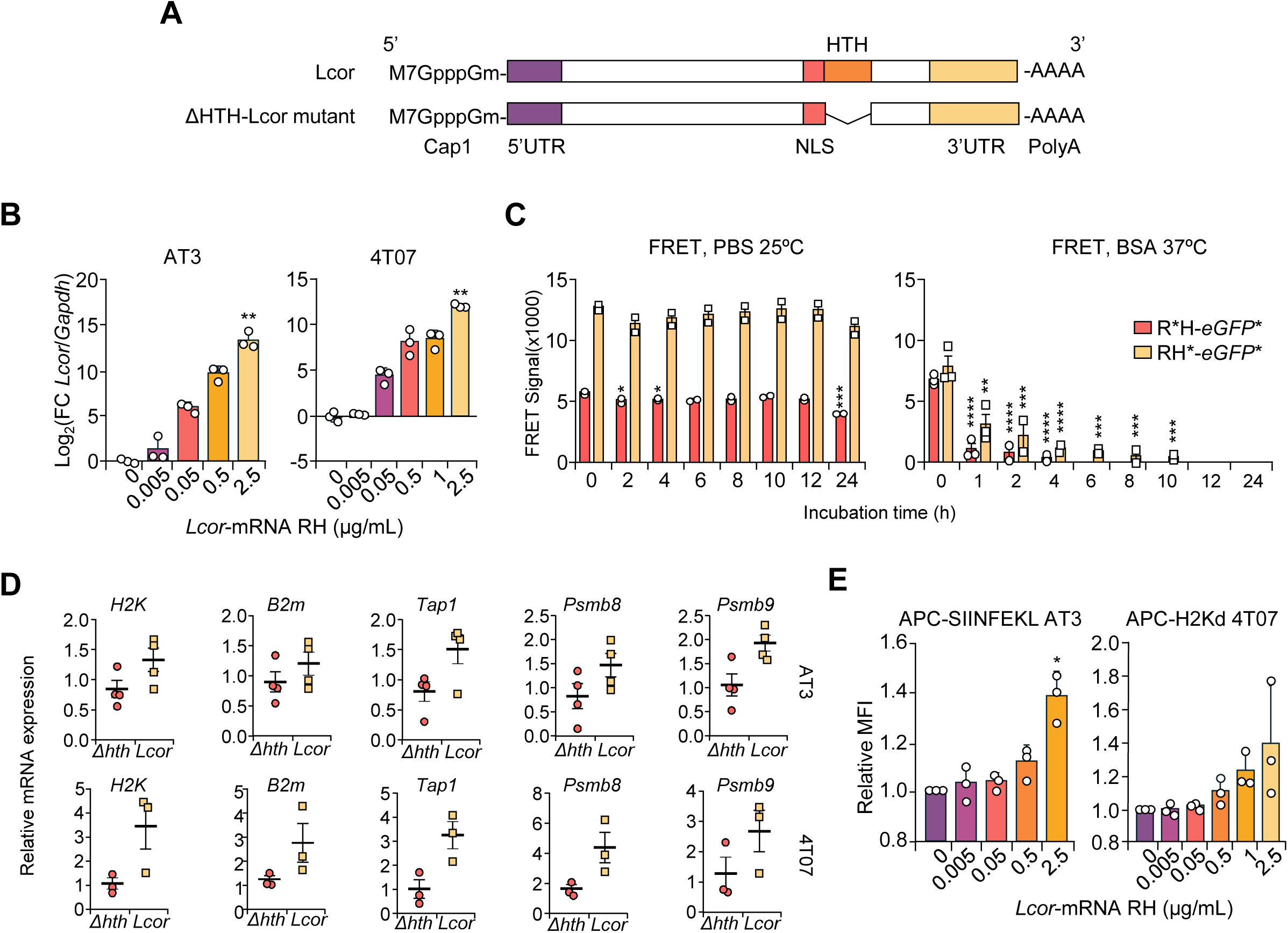
**(a)** Schematic representation of the synthetic mRNA used in this study. The Cap1 structure, 5’ and 3’ untranslated regions, polyA tail, and relevant sequences within *Lcor* mRNA are depicted. **(b)** RT-qPCR analysis of *Lcor* mRNA levels in TNBC AT3 and 4T07 cells treated with the indicated concentration of mRNA-loaded pBAE-NPs for 12 h. Data are represented as Log_2_(Fc) to the normalized expression of non-treated control condition. Expression levels were normalized to *Gapdh*. N = 3 independent experiments; data represent the mean ± S.E.M. **(c)** Stability of the pBAE-NPs over time, measured by FRET efficiency using spectrophotometry. The polymer (R or H) was labeled with Cy5 and the mRNA with Cy3. Stability was assessed at 25°C in PBS or at 37°C in 50 mg/mL of BSA. Data represent mean + S.E.M. from 3 independent experiments. **(d)** Western blot image of TNBC AT3 and 4T07 cells treated with *Lcor* mRNA-loaded pBAE-NPs 6, 12, 24, 36, and 48 h. Detection of ΔHTH-Lcor mutant and Vinculin (loading control). **(e)** Flow cytometry analysis of antigen presentation levels in AT3-OVA and 4T07 cells treated with the indicated dose of *Lcor* mRNA-loaded pBAE-NPs for 12 h. AT3-OVA cells were stained for SIINFEKL-H-2K^b^ complexes and 4T07 for H-2K^d^ to assess peptide-MHC I presentation. N = 3 independent experiments. **(f)** RT-qPCR analysis of APM genes in TNBC AT3 (left) and 4T07 (right) cells treated with 5 µg of the indicated mRNA. Data are represented as Fc to the normalized expression of non-treated control condition. Expression levels were normalized to *Gapdh*. N = 4 independent experiments for AT3 and 3 independent experiments for 4T07; data represent the mean ± S.E.M. Statistical significance; *p<0.05, **p<0.01, ***p<0.001, Kruskal-Wallis H test, followed by Dunn’s post hoc test in B and E, one-way ANOVA test in C, and Mann-Whitney U test in D.

**Figure S3.**
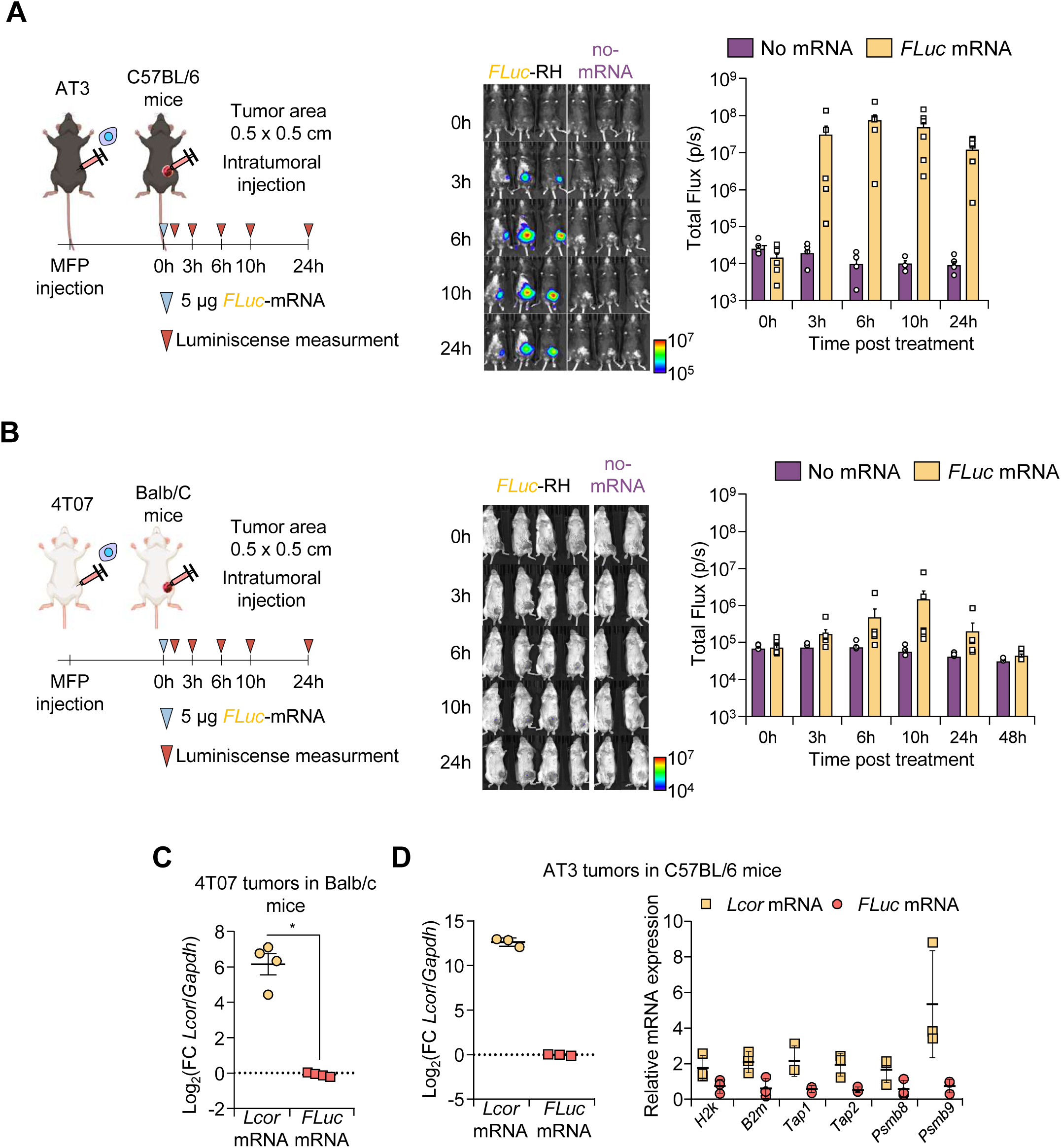
**(a)** *In vivo* protein expression following treatment with FLuc mRNA-loaded pBAE-NPs. AT3 cells were implanted in the mammary fat pad (MFP) of C57BL/6 mice. Once tumors reached 0.5 × 0.5 cm², tumors were treated with 5 µg of FLuc mRNA-loaded pBAE-NPs. Luminescence from translated FLuc protein was measured at 0, 3, 6, 10, and 24 h post-administration using IVIS Spectrum. Representative IVIS images and corresponding quantification are shown. Data represent mean ± S.E.M. N = 3–4 mice per group. **(b)** *In vivo* protein expression following treatment with FLuc mRNA-loaded pBAE-NPs. 4T07 cells were implanted in the MFP of Balb/C mice. After reaching a size of 0.5 × 0.5 cm², tumors were treated with 5 µg of *FLuc* mRNA-loaded pBAE-NPs. Luminescence was measured at 0, 3, 6, 10, and 24 h post-administration using IVIS Spectrum. Representative IVIS images and corresponding quantification are shown. Data represent mean ± S.E.M. N = 3–4 mice per group. **(c)** RT-qPCR analysis of *Lcor* mRNA levels in tumors formed by 4T07 cells implanted in the MFP of BALB/c mice. After reaching a size of 0.5 × 0.5 cm^2^, tumors were intratumorally treated with 5 µg of *Lcor* mRNA-loaded pBAE-NPs and harvested 24 h post-treatment. Data is represented as Log_2_(Fc) to the normalized expression of FLuc mRNA treated control condition. Expression levels were normalized to *Gapdh.* N = X independent experiments; data represent the mean ± S.E.M. **(d)** RT-qPCR analysis of *Lcor* mRNA levels (left) and APM gene expression (right) in tumors formed by AT3 cells implanted in the MFP of C57BL/6 mice. After reaching a size of 0.5 × 0.5 cm^2^, tumors were intratumorally treated with 5 µg of *Lcor* mRNA-loaded pBAE-NPs and harvested 3 h post-treatment. Data are represented as Log_2_(Fc) or Fc to the normalized expression of FLuc mRNA treated control condition. Expression levels were normalized to *Gapdh.* N = 3 independent experiments; data represent the mean ± S.E.M. Statistical significance; *p<0.05, **p<0.01, ***p<0.001, by Mixed-effect analysis in A and B, and Mann-Whitney U test in C and D.

**Figure S4.**
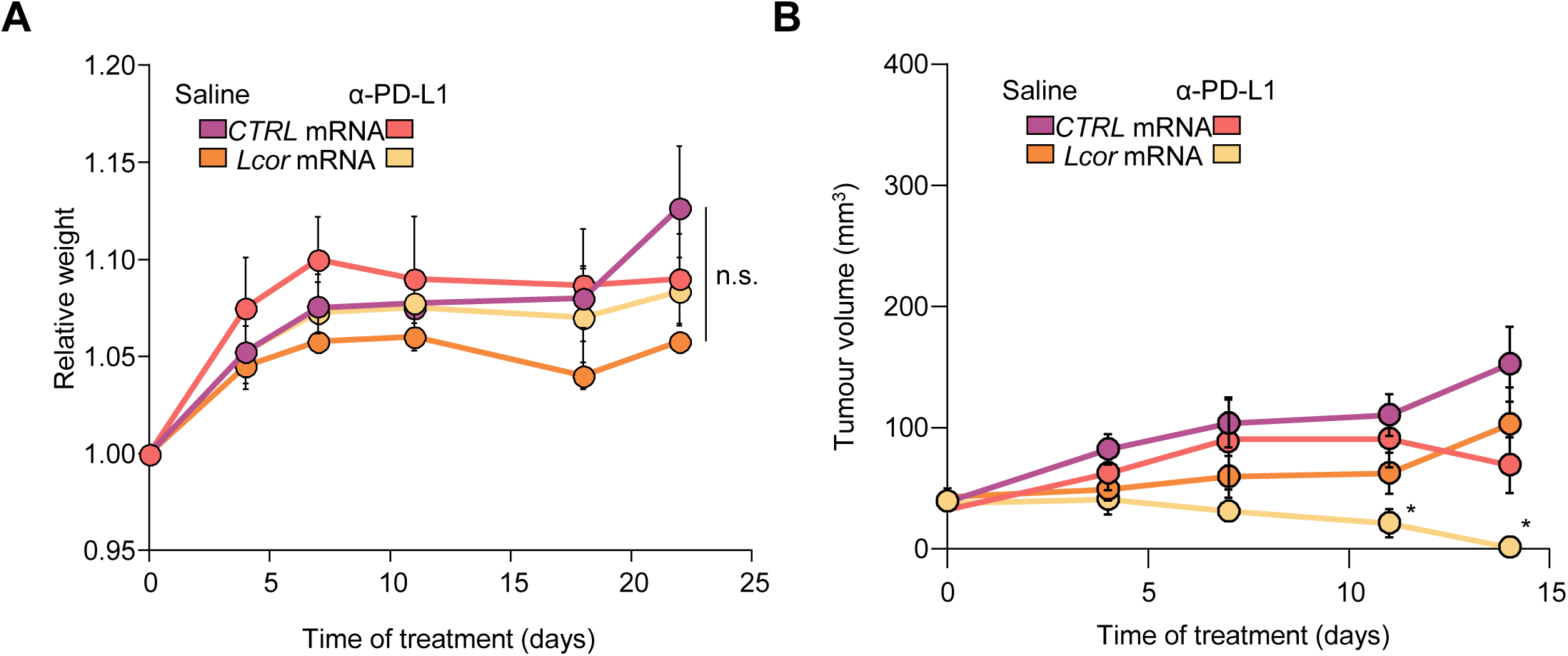
**(a)** Relative weight of mice over time in the specified treatments. **(b)** Tumor volume over time in the specified treatments. Mice were implanted with 4T07 cells in the MFP and tumors treated with 5 µg of mRNA-loaded pBAE-NPs intratumorally twice a week, and 5 mg/kg of anti-PD-L1 intraperitoneally once a week. Data represent mean ± S.E.M. from N = 5–7 mice per group. Statistical significance: *p<0.05, **p<0.01, ***p<0.001, by linear mixed model analysis with Geisser-Greenhouse correction, followed by Tukey’s multiple comparisons test.

**Figure S5.**
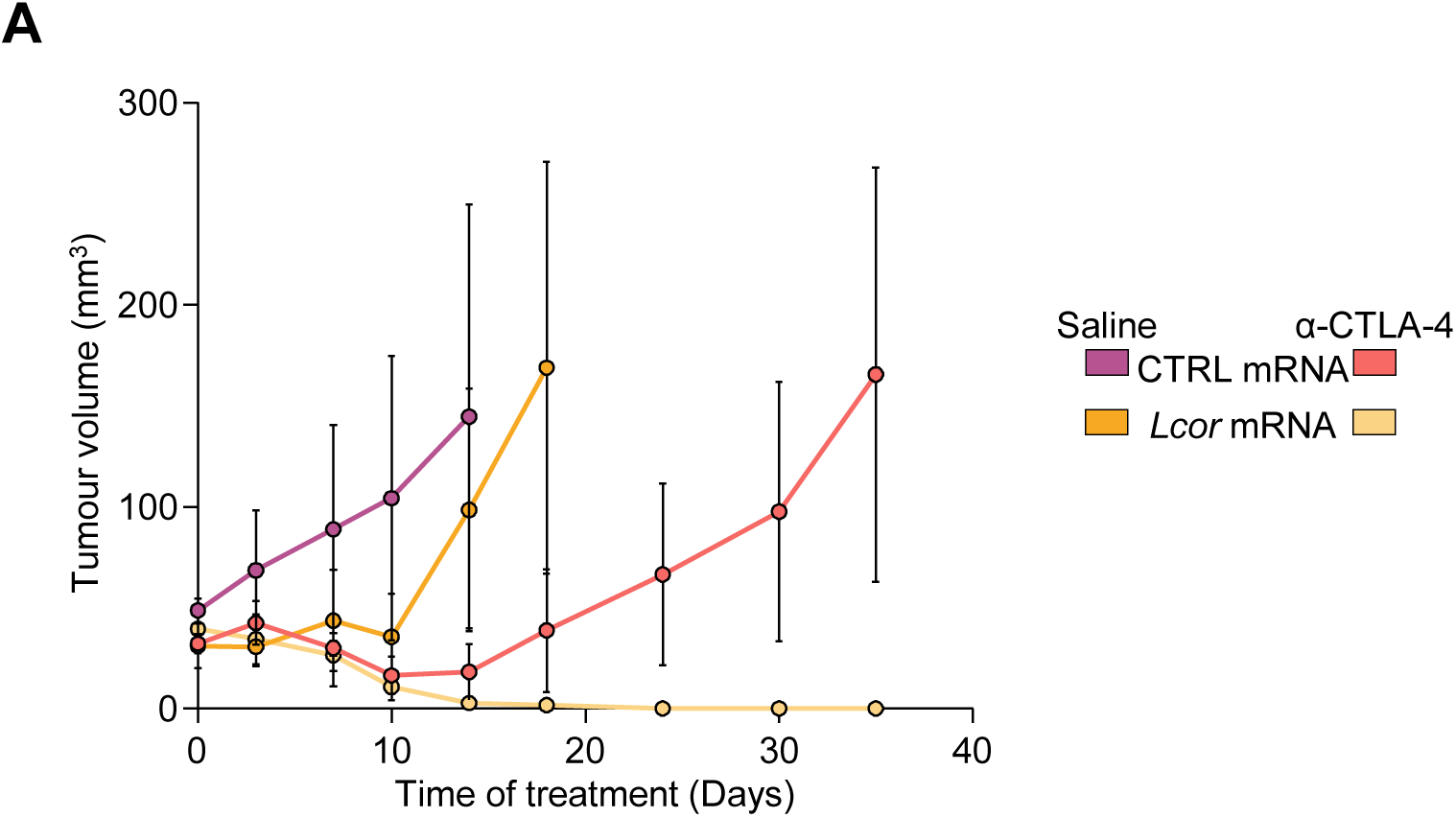
**(a)** Tumor volume over time in the specified treatments. Mice were implanted with 4T07 cells in the MFP and tumors treated with 5 µg of mRNA-loaded pBAE-NPs intratumorally twice a week, and 5 mg/kg of anti-CTLA4 intraperitoneally for the first dose, followed by 0.5 mg/kg for subsequent doses. Data represent mean ± S.E.M. from N = 5–6 mice per group.

**Supplementary Table 1.**
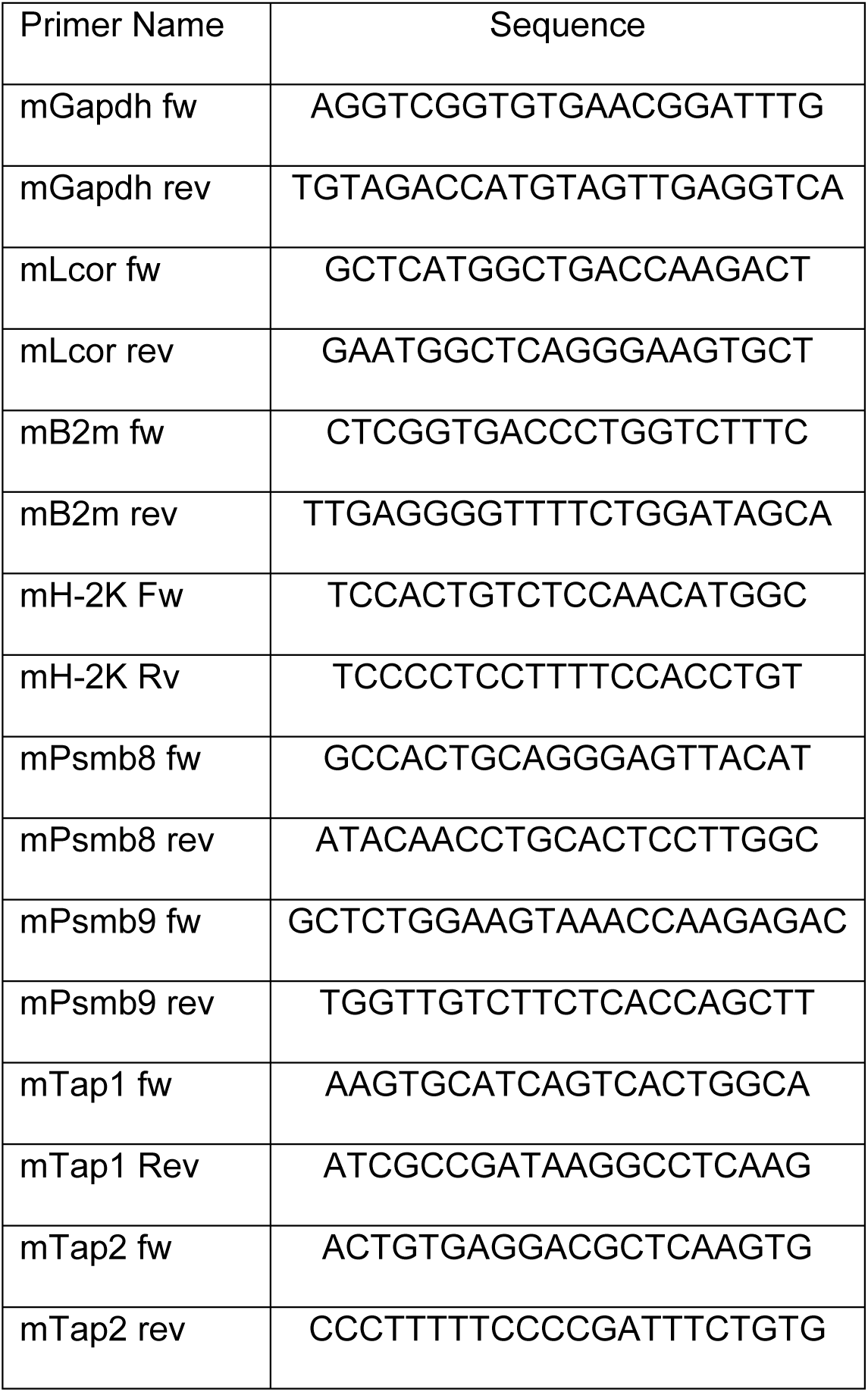

**Supplementary Table 2.**

| Antibody name | Source | ID |
| --- | --- | --- |
| IF and WB: Rb pAB to HA tag | Abcam | AB9110 |
| IF: Goat anti-Rabbit IgG (H+L) cross-Adsorbed secondary antibody AF 555 | Invitrogen | A21428 |
| FC: APC anti-mouse H-2K<sup>d</sup> | BioLegend | 116620 |
| FC: APC Mouse IgG2a, κ Isotype Ctrl | BioLegend | 400220 |
| FC: OVA257-264 (SIINFEKL) peptide bound to H-2Kb | eBioscience | 17-5743-82 |
| FC: Mouse IgG1 kappa Isotype Control (P3.6.2.8.1), APC | eBioscience | 17-4714-82 |
| WB: Goat Anti-Rabbit IgG H&L (HRP) | Abcam | AB6721 |
| In vivo: *inVivo*MAB anti-mouse PD-L1 (B7-H1) | Bioxcell | BE0101 |
| In vivo: *InVivo*MAB anti-mouse CTLA4 (CD152) | Bioxcell | BE0131 |
| In vivo: *InVivo*MAB rat IgG2b isotype control, anti-keyhole limpet hemocyanin | Bioxcell | BE0900 |
| In vivo: *InVivo*MAB polyclonal Syrian hamster IgG | Bioxcell | BE0087 |

